# Rate fluctuations not steps dominate LIP activity during decision-making

**DOI:** 10.1101/249672

**Authors:** Xuelong Zhao, Konrad P. Kording

## Abstract

The idea that lateral intraparietal cortex (LIP) integrates information for and against a decision, is one of the most popular models in neuroscience. However, a recent statistical analysis has suggested that LIP does not integrate information but that individual neurons’ activities jump. The result was based on a model comparison, which is often hard to interpret. There are two worries that can render comparisons problematic. (1) Important aspects of variance are contained in neither model. (2) The analysis is complicated, making it hard to verify. We thus followed up with a simple approach for model comparison: crossvalidation. We find evidence that baseline fluctuations describe much of the variance, which are properly modeled by neither the original paper’s drift-diffusion model, nor simple ramp or step models. Moreover, we find that our straightforward analysis strategy prefers ramping models, both with and without trial-by-trial baseline fluctuations. Our analysis, implementable in a few lines of code, suggests the importance of simple analyses.

---

Most decisions are made under uncertainty and their neural implementation is typically studied using tasks where noisy stimuli need to be integrated over time. For example, noisily moving dots may be shown before asking an animal the question if they predominantly moved left or right in a two alternative forced choice (2AFC, or more precisely a yes-no task) paradigm. This allows asking how information is integrated over time.

Results from such experiments are typically modeled using the drift-diffusion model (DDM) which assumes that neurons linearly integrate weak information for either direction which produces drift of the state while noise adds diffusion. The finding of neurons in LIP that have activities that, once averaged across trials, show ramping activities (Gold and Shadlen, 2007) has been seen as strong evidence for the DDM model. The DDM model has been successful at describing both behavioral and neuronal findings.

A recent paper (Latimer et al., 2015) has cast doubt on this model. They fit a simple model, where the rate jumps up or down at a given point of time with two free parameters, the time of transition and the binary direction. They also fit a very high dimensional DDM model where activity evolves over time and found that the jump model had a better deviance information criterion (DIC). However, there are plenty of reasons to be concerned about the analysis. First, it is too complex for even specialists in the art to replicate. Second, we know that model comparison techniques are notoriously fickle at comparing models that are vastly different. Third, the model did not allow a neurons firing rate to explicitly depend on its own past history, an effect that is known to be large. Lastly, we know from past research that firing rates change across trials Goris et al. (2014), which is modeled by neither model. There is an ongoing debate about the role of LIP in decision making and we think that a simple reanalysis could be illuminating.

Indeed, there are three simple models that one could pose in this space. Either a constant firing rate that is modulated in baseline across trials, a ramp, or a step model, or potentially their combination. In Fig. 2 the models panel shows different variations of the ramp or step model.

There are different strategies to compare models. For example, Akaike Information Criterion (AIC), Bayesian Information Criterion (BIC), and Deviance Information Criterion (DIC) (Burnham and Anderson, 2004), which are all asymptotic approximations that involve complicating assumptions regarding the posterior. A conceptually simpler, alternative approach is cross-validation (CV). CV is a method for estimating a model’s predictive accuracy by dividing the training data into partitions or folds. Each fold is then used iteratively as a test set while the remaining folds are used for training. The model’s average perform is then used to estimate the model’s predictive accuracy. CV boils down to the idea that a good model should generalize well. CV allows us a fresh and above all simple perspective on the question of different models underlying sensory integration in LIP.

Here we use a simple statistical approach for the comparison of three competing models: steps, ramps, and a constant firing rate model. We train the model on data from training time-bins, and test how good each of the models is at predicting spikes in the held-out time-bins. This is a simple metric that allows a meaningful model comparison. Moreover, we compare three simple models, each of which has just a single free parameter to make them more interpretable and easily compared.

## Results

To understand neural activities we first look at some raw traces. We observe that for some neuron/condition pairs, there are trials that elicit many spikes while others elicit few of them (Fig. 1 left). We can quantify this dispersion of spike trains using the Fano factor (Fig. 1). Indeed, Fano factors are often much higher than expected for a Poisson process (FF=1). It seems unlikely that typical models for firing rates would predict such a dispersion of firing rates, after all, the analyzed trials are fixed length. This raises the issue if this effect will introduce biases into a model comparison strategy.

**Figure 1:**
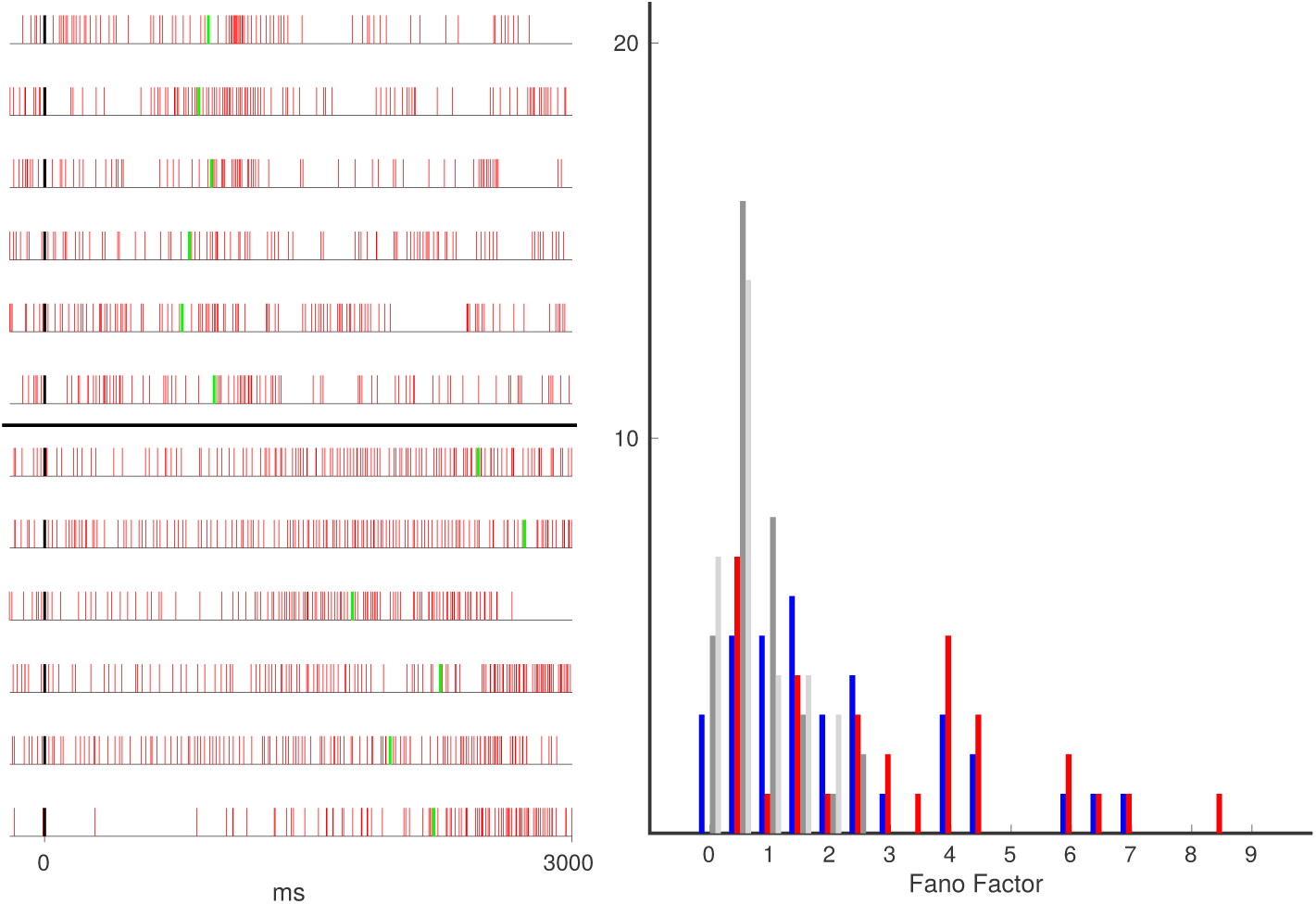
Left: examples of spike trains contributing to high Fano factors (>6) aligned to stimulus onset (black), saccade times are indicated by green lines. Red spike trains are out-RF, horizontal black line denotes spike trains from different conditions. Right: histogram of Fano factors calculated from real and synthetic data belonging to the same neuron, coherence level, and receptive field. Blue: real spike trains, in-RF; red: real spike trains, out-RF; dark-grey: synthetic data, in-RF; light-grey: synthetic data, out-RF.

**Figure 2:**
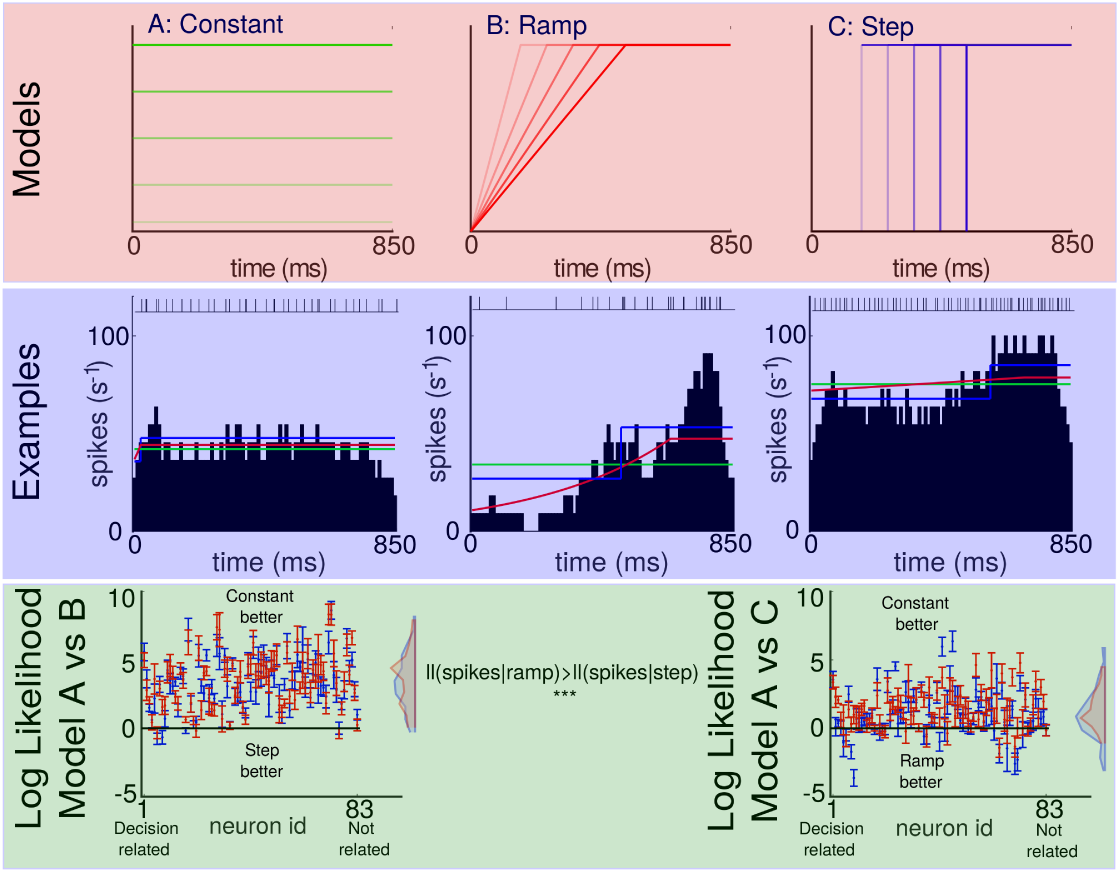
Top: One-parameter variations of the constant, ramp, and step models respectively. Middle: Typical single trial binned (10ms) spike trains of three different neurons with the spike train shown at the top. Fits of the models are superimposed (same colors as in top row). Bottom: Difference of the mean crossvalidated log-likelihoods constant vs step model (left) and constant vs ramp model (right). Red represents model with a self-history covariate, blue represents model without. Error bars are the standard error of the mean across folds.

We thus ask how well the three simple models describe the spikes. We thus analyze the probability each model assigns to the actual spike counts in bins that have not been used to train the model (which includes trial-specific and across-trial parameters). We find that generally the fixed activity model outperforms the other two, and that the ramp model outperforms the step model on individual trials (Fig. 2 Log Likelihood panel). These findings are somewhat surprising given the past literature’s emphasis on ramping models (Hanks et al., 2006; Shadlen and Kiani, 2013; Roitman and Shadlen, 2002; Kiani and Shadlen, 2009; Churchland et al., 2008). It seems like the trial-by-trial variance is dominated by baseline fluctuations, suggesting that the previously observed preference for jump models may simply reflect the fact that jump models can describe more of the firing rate fluctuations. The strong fluctuation of firing rate across trials dominates the variance rendering comparisons problematic.

Clearly, a model in which only the baseline drifts can not adequately describe the observed post stimulus time histograms (PSTHs). In fact, a crossvalidation approach will reject a more complicated model if it does not help at describing individual trials. Interestingly, the predictions from a model that contains both mean fluctuations and also ramping or stepping are worse than those considering only constant fluctuations. As such, our model, that does not regularize, is biased against complex models. Our finding simply mean that in any given trial, we can not meaningfully detect the ramping or stepping. However, we can detect baseline fluctuations, suggesting that they have a dominating effect.

Furthermore, we find that history dependence helps considerably in predicting spikes in time-bins (average log probability change 4.5993±0.5817 across neurons for the constant model). This should not be surprising given previous studies in this space (Pillow et al., 2008; Paninski et al., 2007; Truccolo et al., 2011). History dependence, somewhat to our surprise, seems to have relatively little influence on the preference for ramping, stepping and constant models.

To address the ongoing controversy in the field, we compared, in a conservative manner, different types of models for LIP neurons on individual trials. We found the constant model to work best, but the ramp to somewhat outperform the step model. The strong performance of the constant model is probably due to known massive fluctuations baseline firing rates (Fig. 1) probably due to changes in excitability(Goris et al., 2014). The strength of such an effect makes the comparison between step and ramp models (Latimer et al., 2015) problematic. The jump model has some of the properties of the constant model, if it jumps in the beginning it adds a constant to excitability. Alternatively it can jump into the opposite direction. This apparent modulation of average firing rate is something that the ramp model simply can not achieve early in the trial. It is nearly impossible to interpret the fits of models to data if none of them properly does justice to the data(Kording et al., 2017).

Model comparison is complicated and we have purposefully used one of the most direct approaches. The standard comparison metrics, e.g. AIC, BIC, and DIC are complicated, based on a host of assumptions, and hard to interpret in situations where the real model is not part of the considered set. By restricting all three models of interest to only one degree of freedom, and simply evaluating cross-validated prediction, we used a simple but easy to verify and understand approach. At some level, our paper tells a cautionary tale - if the comparison is too opaque, then details in the approach may decide which model wins.

Obviously one could build better models of neural spiking. We were able to readily consider spike history but that strongly affects log likelihoods but only weakly the distribution of relative log likelihoods between the models. One could also model motor planning, transient responses to fixation, stimulus onset and stimulus offset, but it is unclear which specification we should prefer. Furthermore, the ramp and step models use regressors that are restricted to between 0 and 1, hence we assume they are scaled by the same estimated weight across trials for any particular neuron due to a common covariate weighting. This is ameliorated to an extent by separating integration models related to in-RF and out-RF saccades into separate covariates. The upshot is that there are countless specifications. Which one should be used to ask the question of how the brain makes decisions is unclear.

When the finding that an integration model did worse than a jump model at explaining activity in LIP (Latimer et al., 2015) were published, many commentators where quick to dismiss the drift diffusion model of decision making. However, a close reading of the paper made clear that the comparison was very complicated. Given the possibility of coding bugs in large academic software (Axelrod, 2014), complex analyses are always a worry and even if correct they are hard to replicate. Here we used a much simpler strategy and obtained considerably different answers. This may suggest that we do not do justice to the problem by not fully modeling the drift-diffusion process. Alternatively, it may instead suggest that the original paper does not take into account the constant trial-by-trial changes in baseline. In that sense, we want to simply emphasize that the field needs to use a breadth of model comparison strategies. If the effects are not sufficiently obvious, we should expect that any of the many defensible strategies for comparison will provide materially different answers and we still do not know how we should feel about the drift diffusion model of LIP activity. But lastly and maybe most importantly, our paper tells a cautionary tale about exploratory data analysis. In fact, it seems that trial by trial fluctuations of firing rates can be seen by the naked eye in Figure 2 of Latimer et al. (2015). Omitting any such effect can dramatically affect the results of model comparison.

## Methods

We ask whether a ramp, step, or constant model is most appropriate to describe neuronal spike trains across individual trials during evidence integration in decision-making tasks. To test our hypotheses regarding integration activity during a decision-making task we built ramp, step, or constant firing rate models that include spike history to compare the likelihoods of each model in describing observed spike data.

### Data Analysis

The dataset we are using for our analysis is from (Roitman and Shadlen, 2002) and is publicly available at https://www.shadlenlab.columbia.edu/resources/RoitmanDataCode.html. We have split the data into fixed duration (FD) and reaction-time (RT) trials. For FD trials integration activity is assumed to begin from 200 ms after stimulus onset and terminates at motion offset. For RT trials integration activity is assumed to begin from 200 ms after stimulus onset and terminates 50 ms before saccade initiation, following (Bollimunta et al., 2012), where we use target acquisition time as a proxy for saccade initiation. This duration is used because neuronal activity at the beginning of a trial contains non-integration related activity and the duration leading up to saccadic activity contains signals related to motor planning and preparatory activity (Roitman and Shadlen, 2002).

We analyze only correct trials, wherein the monkey performs fixation appropriately and chooses the correct target. The trials included both in receptive field (RF) and out of RF trials. If a trial contained no spike trains in the duration of interest then it was excluded.

As in (Latimer et al., 2015), we use the same *d’* analysis to order cells with respect to choice selectivity in the 200-700 ms period after dot motion onset for FD trials and 200 ms after motion onset to 50 ms before saccade initiation. The *d*’ measure for a single cell is

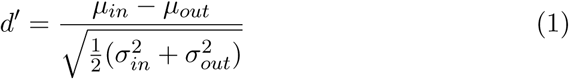

where *μ_in_* and *μ_out_* are mean spike counts on in-RF and out-RF trials respectively. Likewise, 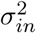 and 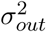 represent variances. Unlike in (Latimer et al., 2015) we analyze all the cells and ordered them using their *d′* values as a proxy for how decision-related the cells were.

The duration between 200 ms after stimulus-on and stimulus-off for each trial was then partitioned into 10ms bins. This allowed a sufficient granularity to distinguish ramping versus stepping activity on the individual neuron level while maintaining computational tractability.

### Generalized Linear Model

We developed a generalized linear model (GLM) approach to estimate the likelihood of ramp, step, or constant models for describing spike trains from neurons on individual trials. Our model describes the probability of a spike rate in a given timebin conditioned on a set of parameters and variables. The time-varying spike rate is given by

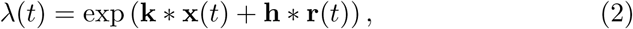

where **k** are kernels for the input variables x of our model and **h** represents the post-spike history filter for the spike history **r**. We used the neuroGLM package (https://github.com/pillowlab/neuroGLM) to find the maximum likelihood estimate of the model parameters for each given trial under an exponential-response GLM model. The model parameters *θ* = **k**, **h** are regularized by ridge regression. We found that the ridge regression constant did not meaningfully affect our results across a range of *ρ* ∈ [0,100], hence we chose *ρ* = 1.0 as our penalty.

We randomly held-out 10% of timebins from each trial during the fitting procedure and validated on the held-out bins during each cross-validation fold. We maximize the log-likelihood of the firing rate of held-out bins from each trial for a given neuron,

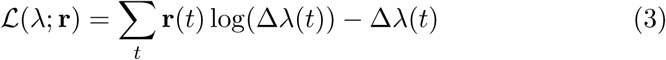

where ∆ is 10 ms.

### Covariates and Integration Models

The design of our GLM model included the binned spike train, a 200 ms spike history filter consisting of raised, log cosines delayed by one timebin with respect to the target spike train, an integration model for in-RF rials, the same type of integration model for out-RF trials, and a constant column as the regressors of interest. The binned spike train and spike history only included spikes from the duration of interest.

We used three types of integration models as possible regressors in our design matrix. The ramp model always begins 200 ms after stimulus onset, and terminated at a chosen timebin, *t_i_*. This implicitly sets the slope of the ramping model, which can be either negative or positive.

For the step model, the chosen timebin, *t_i_*, represented the timebin during which the neuron stepped to a different firing rate. This could either be a step up to a higher firing rate or a step down to a lower firing rate.

The constant rate model takes a constant value which can be chosen from 0 – 100. This is representative of plausible LIP neuronal firing rates.

Both ramp and step models varied from 0 to ±1. The constant integration model ranged from 0 – 100 for each trial. We created separate covariates for integration models in the case of in-RF and out-RF trials.

For all three models, it is possible on each trial to not have a change in the integration model. That is, the ramp or step model is flat during the trial. Also, note that the model covariates are then exponentiated by the generalized linear model that we used here; in fact this alternative model has been posited previously in the literature (Churchland et al., 2011).

### Coordinate Ascent

For a given type of integration model, our goal was to determine the particular variation of the integration model that maximized the log-likelihood of the GLM’s predictions on held-out bins. For all three sensorial integration models we assume evidence integration begins 200 ms after stimulus onset and ends at stimulus offset for FD trials and 50 ms before saccade initiation for RT trials. Hence we restrict our analysis to this duration.

We used coordinate ascent to find the best fit of integration models and weights for the GLM. This is achieved by alternating between fitting the GLM’s covariate weights and finding the best set of timebin *t_i_*’s for trials associated with each neuron. To find the best *t_i_*’s we initialize by setting them randomly, do the first iteration fit of the covariate weights, then fix these weights and scan across all *t_i_*’s for all trials and find the maximum likelihood estimate for each trial. Specifically, in the case of ramps we’ve assumed that integration begins from the first timebin and *t_i_* represents the timebin that the integration ramp hits a decision boundary and terminates. For steps, *t_i_* represents the timebin during which there is a transition to a different firing rate. Finally, for the constant rate model, we do not iterate through the timebins, instead we search across a range of values (0 – 100) and find the best fitting value for each trial.

### Model Comparison

Model comparison is done by finding the mean and variance of log-likelihoods across all *k* folds (*k* = 10) of cross-validation for each integration model type. We then plot the difference of means with standard error mean (sem) between different integration model types as shown in Fig. 2. This comparison is fair as we have only exactly one free parameter for each model type.

Note that Latimer et al. (2015) used latent variables and hence required integration over the latent variable to compute the probability of spike train data, which we have not in this current study.

Additionally, here we have used cross-validation for model selection though it is possible that are situations where it may not be superior to Bayesian approaches to model selection (Shao, 1993).

### Fano Factors

We also study the Fano Factor of spike trains that are from the same neuron, coherence level, and receptive field (in/out). We define the Fano factor as

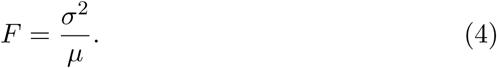

Conditions that contain less than 5 trials were excluded. We then compared this against synthetic Fano factors that we would expect if we assumed the same number of spike trains with the same mean spike rate for each condition were drawn from a Poisson process.

